# SARS-CoV-2 RECoVERY: a multi-platform open-source bioinformatic pipeline for the automatic construction and analysis of SARS-CoV-2 genomes from NGS sequencing data

**DOI:** 10.1101/2021.01.16.425365

**Authors:** Luca De Sabato, Gabriele Vaccari, Arnold Knijn, Giovanni Ianiro, Ilaria Di Bartolo, Stefano Morabito

**Author notes:** Correspondence: Gabriele Vaccari, Emerging Zoonoses Unit, Department of Food safety, Nutrition and Veterinary public health, Istituto Superiore di Sanità, Viale Regina Elena, 299, 00161 Rome Italy.

## Abstract

**Background:** Since its first appearance in December 2019, the novel *Severe Acute Respiratory Syndrome Coronavirus type 2* (SARS-CoV-2), spread worldwide causing an increasing number of cases and deaths (35,537,491 and 1,042,798, respectively at the time of writing, https://covid19.who.int). Similarly, the number of complete viral genome sequences produced by Next Generation Sequencing (NGS), increased exponentially. NGS enables a rapid accumulation of a large number of sequences. However, bioinformatics analyses are critical and require combined approaches for data analysis, which can be challenging for non-bioinformaticians.

**Results:** A user-friendly and sequencing platform-independent bioinformatics pipeline, named SARS-CoV-2 RECoVERY (REconstruction of CoronaVirus gEnomes & Rapid analYsis) has been developed to build SARS-CoV-2 complete genomes from raw sequencing reads and to investigate variants. The genomes built by SARS-CoV-2 RECoVERY were compared with those obtained using other software available and revealed comparable or better performances of SARS–CoV2 RECoVERY. Depending on the number of reads, the complete genome reconstruction and variants analysis can be achieved in less than one hour. The pipeline was implemented in the multi-usage open-source Galaxy platform allowing an easy access to the software and providing computational and storage resources to the community.

**Conclusions:** SARS-CoV-2 RECoVERY is a piece of software destined to the scientific community working on SARS-CoV-2 phylogeny and molecular characterisation, providing a performant tool for the complete reconstruction and variants’ analysis of the viral genome. Additionally, the simple software interface and the ability to use it through a Galaxy instance without the need to implement computing and storage infrastructures, make SARS-CoV-2 RECoVERY a resource also for virologists with little or no bioinformatics skills.

**Availability and implementation:** The pipeline SARS-CoV-2 RECoVERY (REconstruction of COronaVirus gEnomes & Rapid analYsis) is implemented in the Galaxy instance ARIES (https://aries.iss.it).

## Introduction

In December 2019, a novel coronavirus was reported in patients with pneumonia infections in Wuhan, China (Zhu et al., 2020). The novel coronavirus, *Severe Acute Respiratory Syndrome Coronavirus type 2* (SARS-CoV-2), and the related disease, *Coronavirus Disease 2019* (COVID-19) (Gorbalenya et al., 2020), spread rapidly culminating in the WHO declaration of the pandemic state on March 2020, which is still ongoing.

During the pandemic outbreak, NGS technologies enabled complete genome sequencing of thousands of viral strains worldwide and the assessment of temporal and geographical virus spreading (e.g., EpiCOV/GISAID: https://www.gisaid.org).

The NGS technologies produce millions of sequences, however, the manipulation and processing of files can be challenging due to files’ size and can be affected by the lack of bioinformatic skills. The different sequencing standards available (Ambardar et al., 2016) (e.g. Illumina, Ion Torrent, Nanopore) are supported by platforms developed by the companies and made available for the users to a certain extent. On the other hand, the scientific community frequently performs analysis of sequencing data, through commercial software, requiring payment of license keys, or in-house command-line-based pipelines, which demand the availability of bioinformatic skills. In this study, with the intention to provide an all-in-one tool aimed at SARS-CoV2 genomes reconstruction and analysis we concatenated common command-line-based tool into a pipeline, named SARS-CoV-2 RECoVERY (REconstruction of COronaVirus gEnomes & Rapid analYsis), implemented on the multi-usage open-source Galaxy instance ARIES (https://aries.iss.it), dedicated to public health microbiology (Knijn et al., 2020).

## Methods

### Overview

The SARS-CoV-2 RECoVERY consists of six steps: (1) read quality analysis and trimming, (2) subtraction of human sequences, (3) reads alignment and reference mapping against the SARS-CoV-2 reference sequence, (4) variant calling, (5) consensus sequence calling, (6) ORFs identification and variant annotation.

### Building databases

The GenBank file of the reference genome of SARS-CoV-2 (isolate Wuhan-Hu-1; Accession number: NC045512.2) was used to build two databases: a fasta format file containing the complete virus genome used as reference and a database containing the Open Reading Frames (ORF) annotation by the SnpEff tool (Cingolani et al., 2012) in gbk format.

### Read quality analysis and trimming

The reads imported in fastq format are trimmed with the Trimmomatic tool (Bolger et al., 2014) to remove the low-quality bases (or N bases) from both terminus of each read and to exclude reads shorter than 30 base pairs (bp).

### Subtraction of human sequences

Trimmed reads are mapped using Bowtie2 software (Langmead et al., 2012) onto the reference human genome downloaded by “The Genome Reference Consortium” database (https://www.ncbi.nlm.nih.gov/grc) to remove the human genomic sequences

### Genome reconstruction

The recovered unaligned reads are mapped onto the reference sequence of SARS-CoV-2 using the software Bowtie2, for Illumina and Ion Torrent reads, and Minimap2 (Li, 2018) for Nanopore reads.

The resulting BAM file is processed using the iVar consensus caller (Grubaugh et al., 2019) with the following options: minimum quality score threshold to count base 20, minimum frequency threshold 0.6, minimum depth to call consensus 30x.

### Coverage analysis

The coverage analysis and nucleotide distribution are performed using the tool Qualimap 2 (Okonechnikov et al., 2016).

### ORF annotation

Annotation is performed with the BLASTn tool (Megablast) using the SARS-CoV-2 reference ORFs (Open Reading Frame). Because of the high nucleotide identities among SARS-CoV-2 strains, >99% nucleotide identity has been set as a requirement for the ORFs annotation. The parameters used for the alignments are: 1 as maximum number of hits, 80% identity cut-off, and 80% Minimum query coverage per High-scoring Segment Pair (HSP). The output table is converted in a multi-fasta file containing the ORFs identified.

### Variant calling and annotation

The variant calling is carried out with the iVar variant caller (Grubaugh et al., 2019) using the BAM file from the mapping of the cleaned sequencing reads onto the reference sequence of SARS-CoV-2 with the following parameters: minimum quality (Default: 20) and minimum frequency (modified: 0.3).

The SnpEff tool (Cingolani et al., 2012) is eventually used for the variants’ annotation, using the reference genome of SARS-CoV-2 and the iVar output (tsv) converted in vcf file format.

For each of the variants identified, the output consists of: the nucleotide of the reference at each position and the alternative sequence, the codon of the reference and the alternative codon, the nucleotide translation and the information about the mutation (synonymous, missense plus deletions).

### Performance of the pipeline in comparison with other software

One hundred NGS raw data from Illumina, 100 from Nanopore and 50 from Ion Torrent platforms, were downloaded from the NCBI database Sequence Read Archive (SRA). The SARS-CoV-2 genomes from the Ion Torrent and Illumina raw data were built using the pipeline from this study, the CLC Genomics Workbench Ver. 9.5 (Qiagen, Milano, Italy) and the online tool Genome Detective Virus Tool (Vilsker et al., 2019). The Nanopore raw data were analysed only by Genome Detective and our pipeline, since CLC does not accept long reads as input. Finally, the genomes reconstructed from each SRA using the different software, were compared to the corresponding GISAID sequence used as reference. We recorded differences between reconstructed genomes in terms of length difference in comparison with the GISAID reference sequences and number of different nucleotides called, calculated by arithmetic mean.

## Results and Discussion

In this study, we describe the development of a pipeline for the construction and analysis of SARS-CoV-2 genomes and the comparison of the results with those obtained by CLC Genomics Workbench 9.5, Genome Detective Virus Tool using the GISAID sequences as a reference. The SRA used for the analyses were obtained using Illumina, Ion Torrent and Nanopore as sequencing standards and corresponded to the GISAID entries used as reference and downloaded from NCBI database. Most of the genomes built using our pipeline were longer (34 nucleotides on average) than the corresponding GISAID references and those built by CLC and Genome Detective for all the sequencing standards (Table 1). In detail, 96% (48/50) of Ion Torrent, 73% (73/100) of Illumina and 97% (97/100) of Nanopore raw reads produced longer genomes when our pipeline was used. Additionally, these genomes presented less nucleotide differences (n≤7, mean) than the genomes built with other software when compared to the GISAID sequence used as reference.

This finding is of particular interest as such differences may include either incorrect or missing nucleotide assignment, which would hamper the studies on SARS-CoV2 evolution and distribution, since the mutations described so far in SARS-CoV-2 genomes are mainly single point mutations. Since the discovery of the SARS-CoV-2 and the first complete genome sequencing (Wu et al., 2020), 470,276 genomes have been submitted to the GISAID database allowing the prompt identification of mutations, together with geographical and temporal mapping of the circulating strains. In addition, 5 major lineages have been reported worldwide (A, B, B.1, B.1.1, B.1.177), defined by SNP differences. Recently, a novel lineage (named B.1.1.7) was detected within the COVID-19 Genomics United Kingdom (COG-UK) Consortium and characterized by 14 non-synonymous mutations and 3 deletions (Rambaut et al., 2020). To test our pipeline, 6 SRA from Ion Torrent, Nanopore, Illumina technologies were tested and the B.1.1.7 complete genomes were successfully reconstructed, reporting all the deletions and the mutations in the annotation table.

Besides Whole Genome Sequencing, bioinformatics analyses are pivotal to obtain the final results. The pipeline developed in this study is publicly accessible through the Galaxy instance ARIES (https://aries.iss.it) and provides a user-friendly interface, allowing the complete reconstruction of SARS-CoV-2 genomes in 4 to 60 minutes for NGS data composed by 50 thousand to 6 million reads, depending on both the file size and the jobs load on the server. The analyses can be run independently from the users’ hardware and the software can be accessed upon direct registration on the ARIES home page using any browser running on desktop or mobile devices. In addition, ARIES does not request access to the users’ data but is meant to provide a service to the scientific community to boost the knowledge on the evolution of the SARS-CoV-2 in the attempt to favour a global response to this global threat.

**Table 1.** Comparison of GISAID reference genomes with those built by CLC, Genome detective and the SARS-CoV-2 RECoVERY. The consensus sequences built with the different software in the table and used for comparison were obtained using the short reads downloaded from the NCBI SRA corresponding to the GISAID entries, which were used as references.

| Ion Torrent data (n° of analysed runs =50) |  |  |  |  |  |  |
| --- | --- | --- | --- | --- | --- | --- |
|  | Mean difference*<br>in consensus length | Min-Max of<br>difference in<br>consensus length |  | % of consensus<br>sequences longer<br>then GISAID<br>reference | % of consensus<br>sequences with<br>different<br>nucleotide call | Mean** of n° of<br>different nucleotide<br>call |
| CLC | -137 | -544 | +47 | 2 (4%) | 28 (56%) | 4 |
| SARS-CoV-2<br>RECoVERY | +34 | -488 | +106 | 48 (96%) | 48 (93.7%) | 7 |
| Genome<br>Detective | -5172 | -18454 | +11 | 0 (0%) | 49 (99.9%) | 41 |
| Illumina data (n° of analysed runs =100) |  |  |  |  |  |  |
| CLC | -1173 | -8345 | +643 | 18 (18%) | 20 (20%) | 7 |
| SARS-CoV-2<br>RECoVERY | +135 | -1652 | +3379 | 73 (73%) | 52 (52%) | 2 |
| Genome<br>Detective | -167 | -8925 | +1989 | 40 (40%) | 43 (43%) | 16 |
| Nanopore data (n° of analysed runs =100) |  |  |  |  |  |  |
| SARS-CoV-2<br>RECoVERY | +107 | -236 | +1856 | 97 (97%) | 96 (96%) | 3 |
| Genome<br>Detective | +267 | -3816 | +2127 | 91 (91%) | 90 (90%) | 13 |

\* Difference in consensus length: calculated as the arithmetic mean of the differences between the corresponding genome built by each software and the GISAID reference length.
\*\* Different nucleotide call: calculated as the arithmetic mean of the number of different sites (nucleotides) between the GISAID reference and the corresponding genome built by each software.

The simplicity of use and the production of a comprehensive report with all the variants characterized, make this pipeline a valuable tool particularly for scientists with little or no skill in bioinformatic.

## Conclusions

In conclusion, we developed a pipeline for the complete genome reconstruction and analysis of sequence data to help and speed the scientific community in the analysis of SARS-CoV-2 sequencing data. The analyses have been completely automated, and the user interface has been designed to minimize the input from the user in order to provide a support also for the non-bioinformaticians and to enlarge the base of scientists analysing such data.

The release of the software as an open-source pipeline through a Galaxy instance will also allow the scientific community to use this collaborative platform in a reproducible way for the crowdsourcing-based advance of our understanding of this new virus and the different evolutionary scenarios.

## Authors’ contributions

All authors contributed to writing the paper, LDS and GI tested the software, AK and SM developed and designed the software, GV, IDB and SM conceived the project idea and provided advice and assistance throughout the development of the software and the manuscript writing process.

